# CADENCE: Clustering Algorithm - Density-based Exploration and Novelty Clustering with Efficiency

**DOI:** 10.1101/2025.02.24.639863

**Authors:** Lexin Chen, Daniel R. Roe, Ramón Alain Miranda-Quintana

**Affiliations:** Department of Chemistry, University of Florida, Gainesville, Florida 32611, USA; Quantum Theory Project, University of Florida, Gainesville, Florida 32611, USA; Laboratory of Computational Biology, National Heart, Lung, and Blood Institute, National Institutes of Health, Bethesda, Maryland 20892, USA

**Keywords:** algorithms, cluster chemistry, molecular simulation

## Abstract

Unsupervised learning techniques play a pivotal role in unraveling protein folding landscapes, constructing Markov State Models, expediting replica exchange simulations, and discerning drug binding patterns, among other applications. A fundamental challenge in current clustering methods lies in how similarities among objects are accessed. Traditional similarity operations are typically only defined over pairs of objects, and this limitation is at the core of many performance issues. The crux of the problem in this field is that efficient algorithms like *k* -means struggle to distinguish between metastable states effectively. However, more robust methods like density-based clustering demand substantial computational resources. Extended similarity techniques have been proven to swiftly pinpoint high and low-density regions within the data in linear *O(N)* time. This offers a highly convenient means to explore complex conformational landscapes, enabling focused exploration of rare events or identification of the most representative conformations, such as the medoid of the dataset. In this contribution, we aim to bridge this gap by introducing a novel density clustering algorithm to the Molecular Dynamics Analysis with *N* -ary Clustering Ensembles (MDANCE) software package based on *n*-ary similarity framework.

## Introduction

Cluster analysis is a type of unsupervised learning, where data is classified without the use of predefined labels, relying instead on the inherent structure of the data. This method is particularly valuable in analyzing data from molecular dynamics (MD) simulations, where it helps to identify key configurations of a system. It can serve as a foundation for developing Markov State Models, virtual screening, and free energy perturbation calculations. ^1–7^ Non-hierarchical clustering methods are widely used due to their ease of implementation and scalability. Techniques like *k* -means^1,8,9^ and *k* -medoids^10^ are frequently applied. Another method, Radial Threshold Clustering (RTC), was developed by Daura and his team. ^11,12^

Density-Based Spatial Clustering of Applications with Noise (DBSCAN) addresses several shortcomings of the *k* -means algorithm, including its inability to handle noisy data, its limitation to globular clusters, and the need to predefine the number of clusters. ^13^ DBSCAN is a density-based method that uses two key parameters: the radius for forming epsilon neighborhoods and the minimum number of samples required to identify a dense region. Initially, each data point forms its own epsilon neighborhood, which includes all other points within a specified radial distance. A core sample is identified if its epsilon neighborhood contains at least the minimum number of samples. The algorithm then starts with a random core point and expands the cluster by including all points within its epsilon neighborhood, iterating outward until no further points can be added. While this allows DBSCAN to distinguish between high- and low-density regions, it is sensitive to the choice of parameters, particularly epsilon and the minimum number of samples. Additionally, DBSCAN can be prone to forming small clusters and is vulnerable to noise, ^14^ making it computationally expensive, with best-case time complexity of O(*N* log *N*)^13^ and worst-case time complexity of O(*N* ^2^) as every point must be compared to all others to determine its neighbors.^15^ These issues become even more pronounced in high-dimensional datasets. To address these limitations, HDBSCAN (Hierarchical DBSCAN)^16^ was introduced as an extension of DBSCAN, incorporating a hierarchical approach that dynamically adjusts thresholds at each iteration.

HDBSCAN allows for merging or splitting clusters based on their stability, eliminating the reliance on a fixed epsilon parameter and making the method more flexible and robust, particularly in datasets with varying densities or complex structures. This improvement enables HDBSCAN to perform more effectively in challenging clustering scenarios, offering a more adaptive solution compared to DBSCAN. Furthermore, alternative approaches like Density Peaks Clustering (DPC)^17^ have been introduced, which identifies cluster centers as points with significantly higher density compared to their neighbors, without requiring a pre-defined number of clusters. Density peaks are particularly useful in cases with varying densities, as they can adapt to the data’s structure.

We previously released eQual (Extended Quality),^18^ which finds clusters based on a radial threshold. This clustering method is great at finding compact clusters. CADENCE builds on eQual with an addition of outlier expansion. CADENCE has the potential to identify core states for Markov state models (MSMs), which are crucial for understanding complex biological processes such as protein folding and protein-ligand (un)binding. MSMs help in determining the rates between metastable states. Core states represent the most important conformations for describing a system’s dynamics and serve as the input for constructing MSMs. As a density-based clustering method, CADENCE is well-suited for identifying densely populated regions within the phase space, making it useful for building MSMs. One key benefit of using density-based clustering for core state identification is its ability to detect not only metastable states but also states that are less strongly metastable.^19^ Since this applies to most biological systems, CADENCE can effectively identify metastable states of various sizes and densities. This is achieved through its radial threshold search for high-density areas and its nearest neighbor search for lower-density parts of the clusters.

## Theory

The crux of the linearity of the algorithm lies in the *n*-ary similarity framework, where multiple data points can be compared simultaneously. ^20–22^ This advantage has given the extension to more efficient clustering algorithms. ^9,23,24^ The Clustering Algorithm - Density-based Exploration of Nearest Common Environments (CADENCE) is an enhancement of the eQual algorithm. To summarize, eQual begins by selecting seeds for each iteration, then accepts data points within a specified radial distance from the seed to form a cluster. ^18^ The cluster with the highest density and compactness is chosen as the cluster for that iteration, and the process continues until all data points are assigned. A limitation of eQual is its tendency to form primarily globular clusters, which makes it ineffective at identifying regions of varying densities in the data. CADENCE improves upon this by first identifying outliers in the cluster formed by eQual in a given iteration. It then includes any points within a specified distance from these outliers, expanding the cluster until no further points meet the threshold criteria. This iterative process continues until no additional points can be added, after which the algorithm proceeds to the next iteration. n out is the number of outliers to begin the outlier expansion process and this is determined by complimentary similarity. The points with the highest complimentary similarity of the cluster will search for potential outliers to bring to the clusters. Acceptable outliers will be determined from the available points (not a member of any clusters) that are within the expansion threshold to the n out. n point is the number of points that each n out will find in each iteration of the outlier expansion process.

## Materials and Methods

### Molecular Dynamics Simulations

***β*-Heptapeptide** The topology and trajectory files correspond to publicly available data and were assessed through GitHub.^25,26^ The atom selection follows Daura *et al*,^27^ Lys2 to Asp11, with N, C*α*, C, O, and H atoms. The terminal and side chain residues were ignored to minimize noise in the clustering. The single-reference alignment was done after aligning to the 1000^th^ frame.

### Clustering

CADENCE clustering was done using the software package MDANCE, available on GitHub https://github.com/mqcomplab/MDANCE. Both the number of points (n points) and the number of outliers (n out) were thoroughly scanned to study their impact on the final clustering results.

## Results

CADENCE, as a density-based method, overcomes eQual’s limitations and is capable of identifying non-convex, arbitrarily shaped clusters. Whereas eQual specializes in detecting tight and mostly globular-shaped clusters, CADENCE’s flexibility enables it to detect complex, non-convex shapes in the data successfully, as illustrated in Fig. 2. This can be useful when the cluster shape is irregular or the clusters may have varying densities.

**Figure 1.**
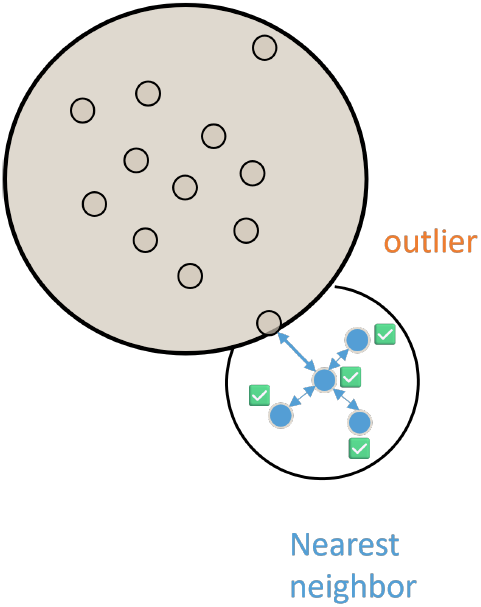
CADENCE schematics.

**Figure 2.**
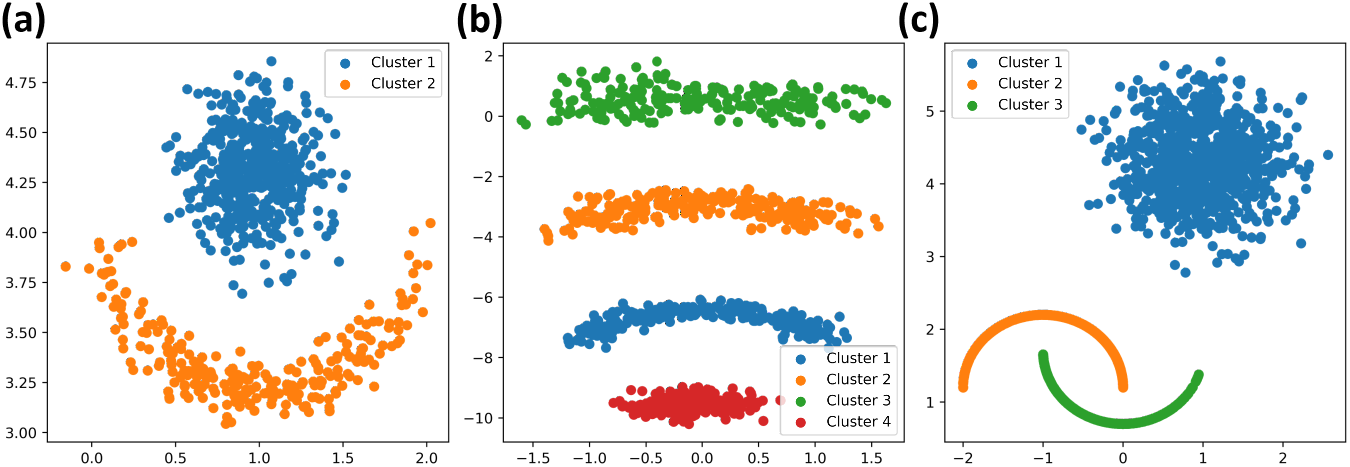
CADENCE clustering on three sample 2D datasets created from scikit-learns

Linear dimensionality reduction methods, such as principal component Analysis (PCA), are effective for visualizing clusters by creating linear combinations of key features. PCA can reduce multi-dimensional data to 2D with the first principal components (PCs) being the eigenvectors with the highest eigenvalues. Nonlinear methods like t-SNE and UMAP, on the other hand, map high-dimensional data to lower dimensions while preserving the relationships and structures based on proximity or similarity between points, making them particularly useful for visualizing complex, non-linear clusters in data. We are looking into all three-dimensional reduction methods.

We compare CADENCE results to eQual’s to see if the arbitrary non-convex shape can be observed. From PCA, tSNE, and UMAP, the general shape of the data is preserved between CADENCE and eQual. In PCA, there are more CADENCE clusters with non-convex shapes. This is further exemplified in t-SNE and UMAP. In eQual, most of the bigger globular sectors consist of one cluster. However, in CADENCE, this is less so the case where a cluster can consist of more than one globular structure, thus highlighting the greater potential discriminative power of CADENCE.

As a further test of CADENCE’s behavior under parameter changes, we sampled through the n out and n point parameters. The criteria to check for cluster quality are the number of resulting clusters, Davies-Bouldin Index, Calinski-Harabasz Index, structures, and population of the top ten clusters.

When sampling varying MSD and n out values (Fig. 5a), there is a peak threshold (resulting in the maximum number of clusters), which is when MSD=1.6. This value is unanimous for the majority of the n out values. Additionally, the peak threshold for CADENCE using n out parameter is aligned with the peak threshold for eQual (MSD=1.7). The shape of the 3D plot is triangular, indicating that MSD has a larger influence on the number of clusters than the number of outliers. Around the peak, more clusters were found when n out values were between 1 to 10 and decreased when greater than 10. When there are more n out there is a greater number of points included in a given cluster, so the number of clusters will be smaller because there are fewer points left to explore, next, when sampling varying MSD and n points values (Fig. 5b), the maximum number of clusters is also between 1.9 to 2.2. However, in this case, the cluster number increases with more points selected. The remarkable stability of the MSD vs n out is a strong indication that CADENCE is capable of.

**Figure 3.**
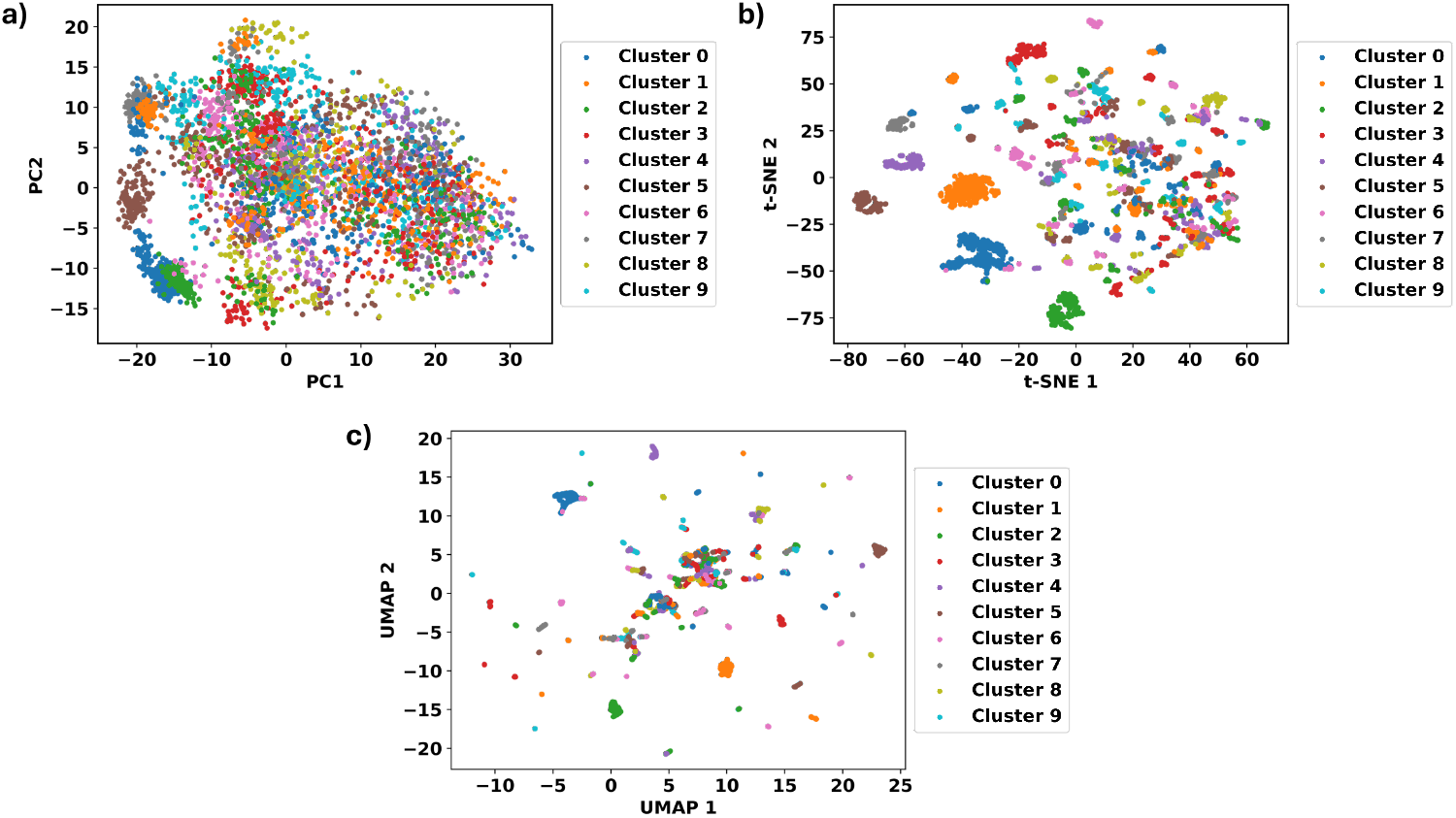
First two components of eQual clustering results at the maximum number of clusters (*k* =155 and MSD=1.7) **(a)** using PCA, **(b)** tSNE, and **(c)** UMAP.

**Figure 4.**
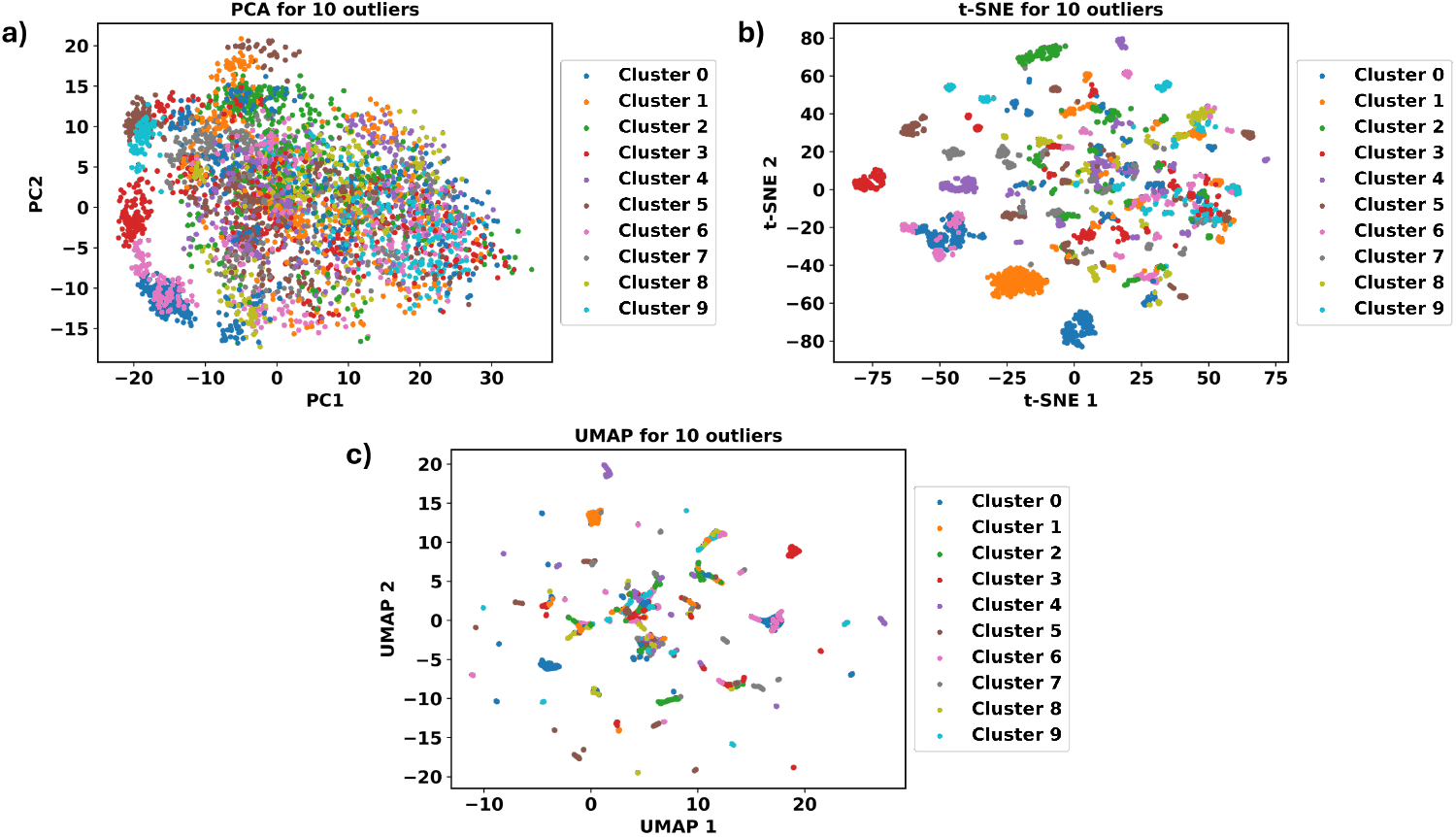
First two components of CADENCE clustering results when the number of outliers is 10 and MSD is 1.6 **(a)** using PCA, **(b)** tSNE, and **(c)** UMAP.

**Figure 5.**
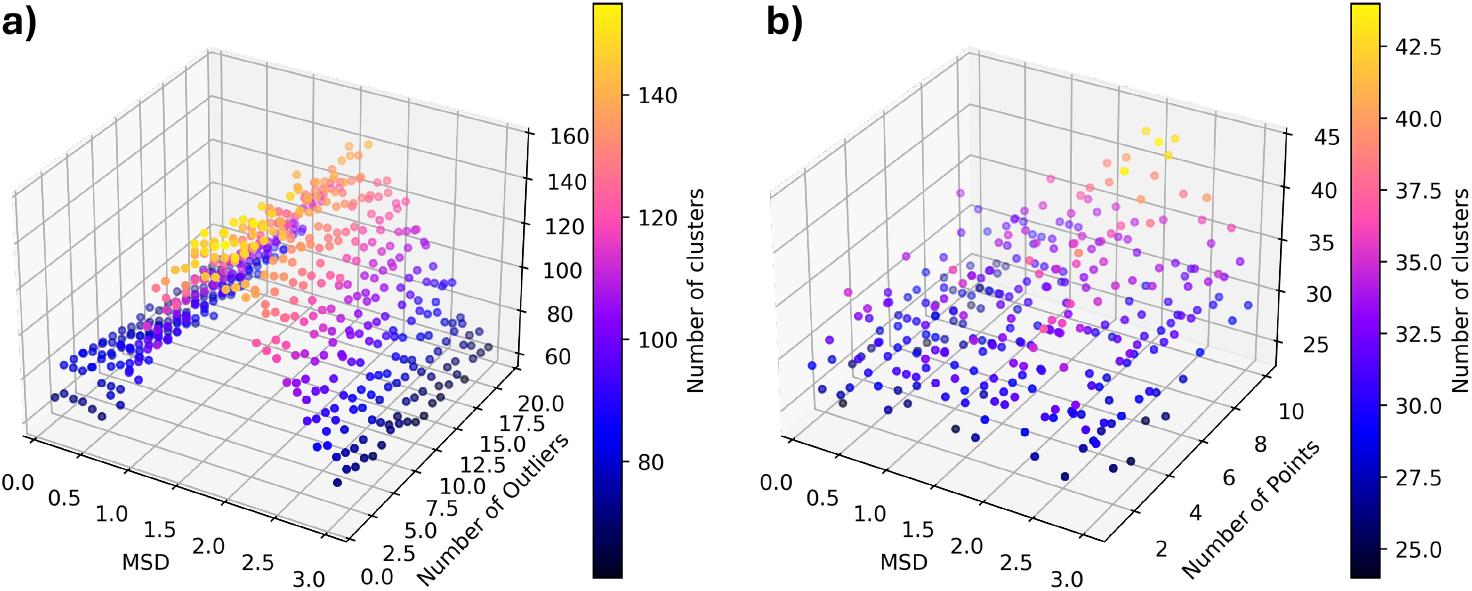
Screening of the number of clusters using two parameters, **(a)** MSD and number of outliers **(b)** MSD and number of points parameters. *x* -axis is the MSD values, *y* -axis is the number of outliers/points, and *z* -axis is the number of clusters.

CADENCE shares a limitation with every single density-based clustering (like DBSCAN or Jarvis-Patrick), in that they need multiple parameters to determine the best clustering conditions. Therefore, two scoring metrics were used to identify the best parameters for the clustering results. Davies-Bouldin Index (DBI) and Calinski-Harabasz Index (CHI) calculate how well-separated and compact the resulting clusters are, respectively. The trend in DBI is that a lower DBI indicates a more well-separated cluster. Fig. 6 shows that lower MSD results in more well-separated clusters and the scores remain constant with varying numbers of outliers. Fig. 6a shows the global minimum of the DBI for constant MSD, for MSD less than 0.5, a higher number of outliers results in a lower DBI, meaning more well-separated clusters. However, a lower number of outliers results in lower DBI for MSD greater than Fig. 6b shows the maximum second derivative for constant MSD, it follows an opposite trend that MSD less than 0.5, a lower number of outliers are indicated as a local minimum or second derivative maximum. While MSD above 2.5, a greater number of outliers results in more local minima. A similar trend is observed in the scoring plot as the number of clusters figures, which is that MSD has a greater effect on CHI and DBI than the number of outliers. This makes sense because the expansion threshold set is constant when it reaches to larger MSD, fewer nearest neighbors will be accepted to the cluster since they might already be accepted in the cluster before expansion. Global min DBI may not be as useful sometimes as it gives a lower number due to the biases in this indicator, but the 2^nd^ derivative maximum aligns better with the highest number of clusters found when MSD is constant. For CHI, a higher value leads to a more compact cluster. The points corresponding to the global maximum align well with the second derivative minimum. When MSD is less than 1.0, it favors a lower number of outliers and it increases when greater than 1.5.

**Figure 6.**
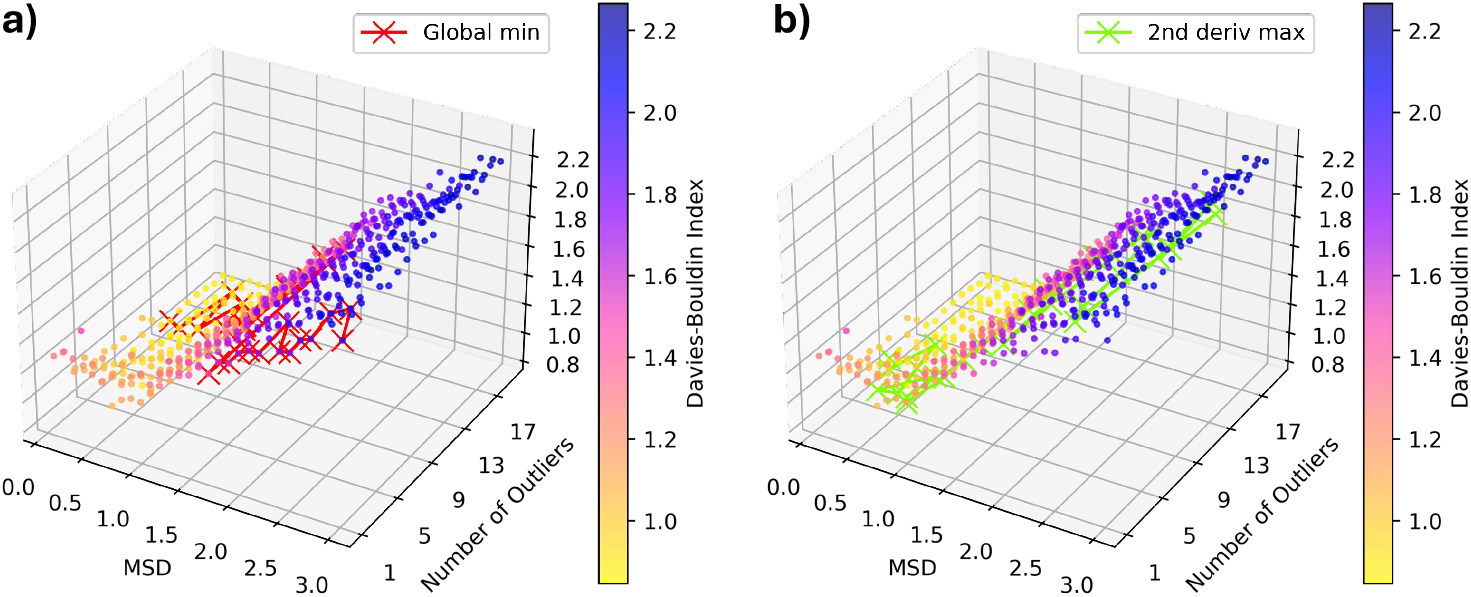
Screening of DBI using two parameters, MSD and the number of outliers. With MSD staying constant, **(a)** marks points with the minimum global DBI value, and **(b)** marks points with the maximum 2nd derivative DBI value.

**Figure 7.**
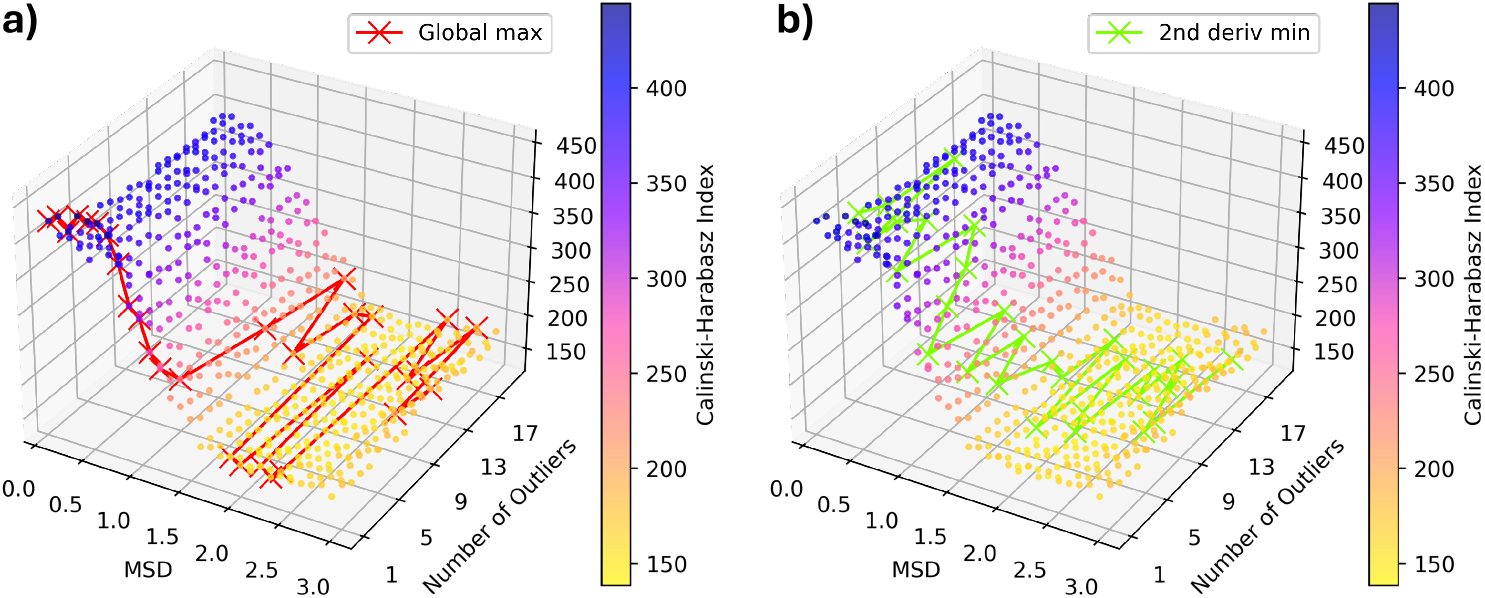
Screening of CHI using two parameters, MSD and the number of outliers. With MSD staying constant, **(a)** marks points with the maximum global CHI value, and **(b)** marks points with the minimum 2nd derivative CHI value.

From the CADENCE populations it can be seen that most of the outlier expansions are concentrated on the biggest cluster. CADENCE clustering has about 7.5% of the frame while eQual has 5.7%, which is perfectly in line with CADENCE giving the eQual clusters a chance to “grow” from their periphery. This can also be seen as the number of outliers (n out) increased, the top cluster had a slightly higher population. This aligns with our method because with more n out looking for nearest neighbors to gather to the cluster, the more likely it is to find points within the expansion threshold to the n out. After cluster 3, the outlier expansion incorporates fewer points, as they are lowly populated and very close to the eQual distribution, so it is further away from the other simulation frames. The population makeup is similar to eQual clustering (Fig. 8) from cluster 4 onward, indicating that those clusters were already quite tight and self-contained from the eQual algorithm.

**Figure 8.**
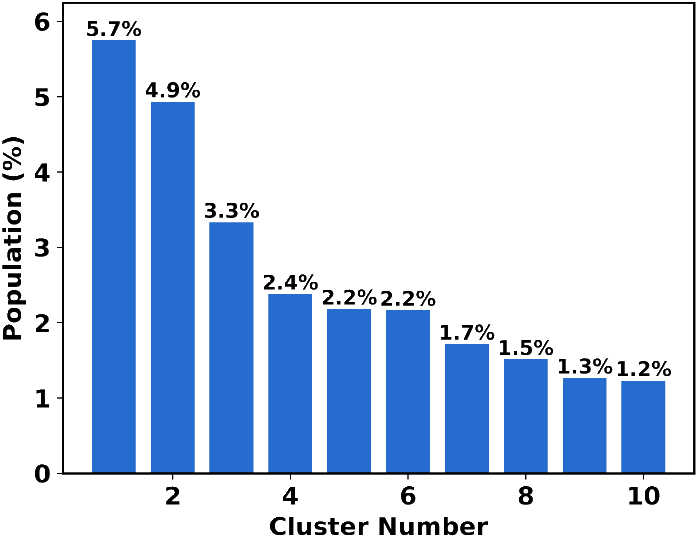
Population of eQual clustering at the maximum number of clusters (*k* =155 and MSD=1.7).

**Figure 9.**
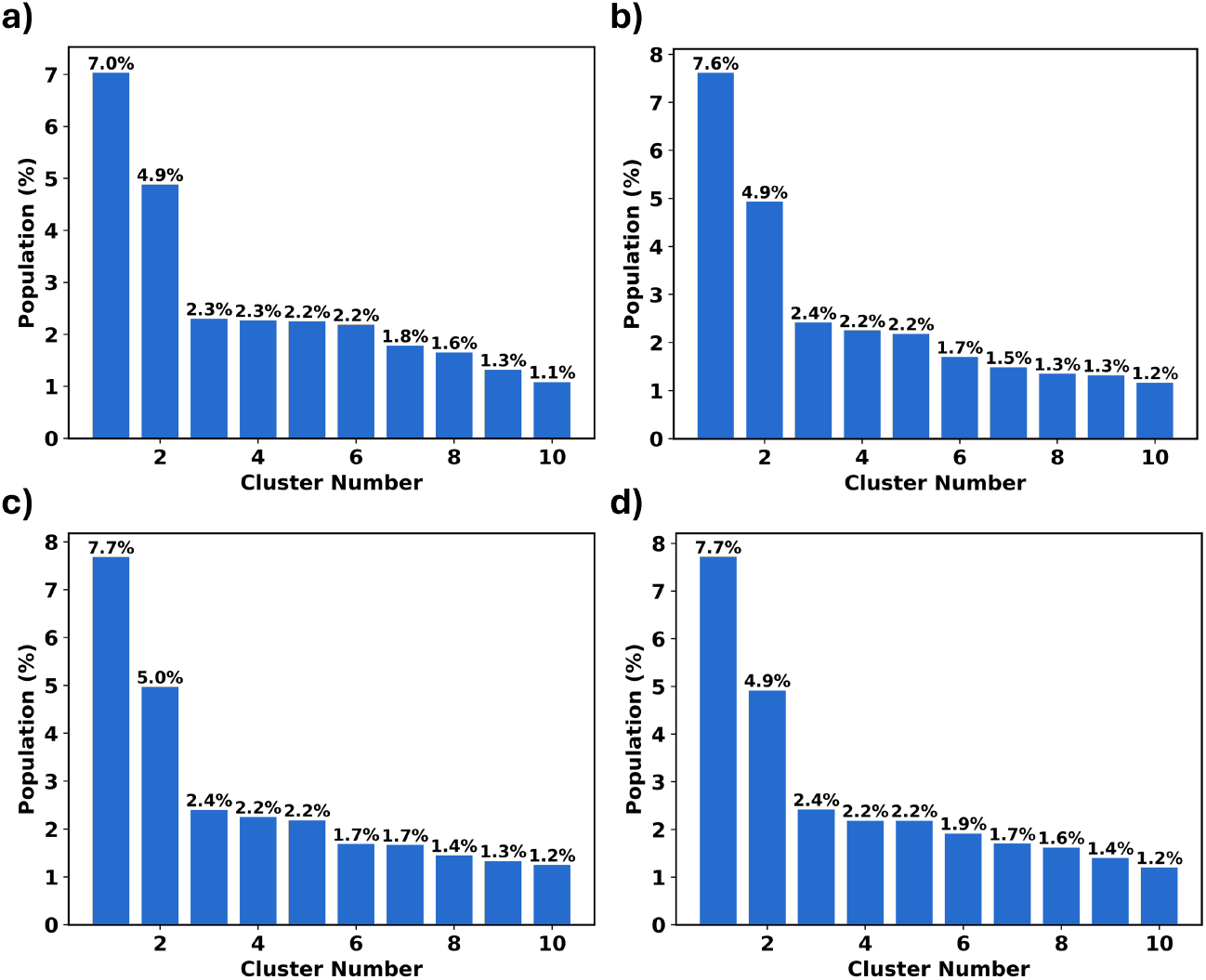
Population of CADENCE clustering results at the maximum number of clusters when MSD is 1.6 and the number of outliers is **(a)** 1, **(b)** 5, **(c)** 10, and **(d)** 19.

Finally, CADENCE was compared to eQual in terms of overlaps of the conformations assigned to each cluster. Figures 11 and 10 illustrate the conformations of the top ten clusters identified at the maximum number of clusters. These conformations align with those reported by Gonzalez-Aleman, ^25^ showcasing the characteristic “U”-, “S”-, “E”-, “3”- and “scarf”-like patterns. Following our previous convention, we show the overlap between cluster conformations (depicted in blue) and the cluster representative (medoid, shown in orange). It is reassuring that CADENCE found similar conformation as eQual while also retaining the tightly compacted cluster nature of the latter. while exploring potential outliers to add to the clusters.

**Figure 10.**
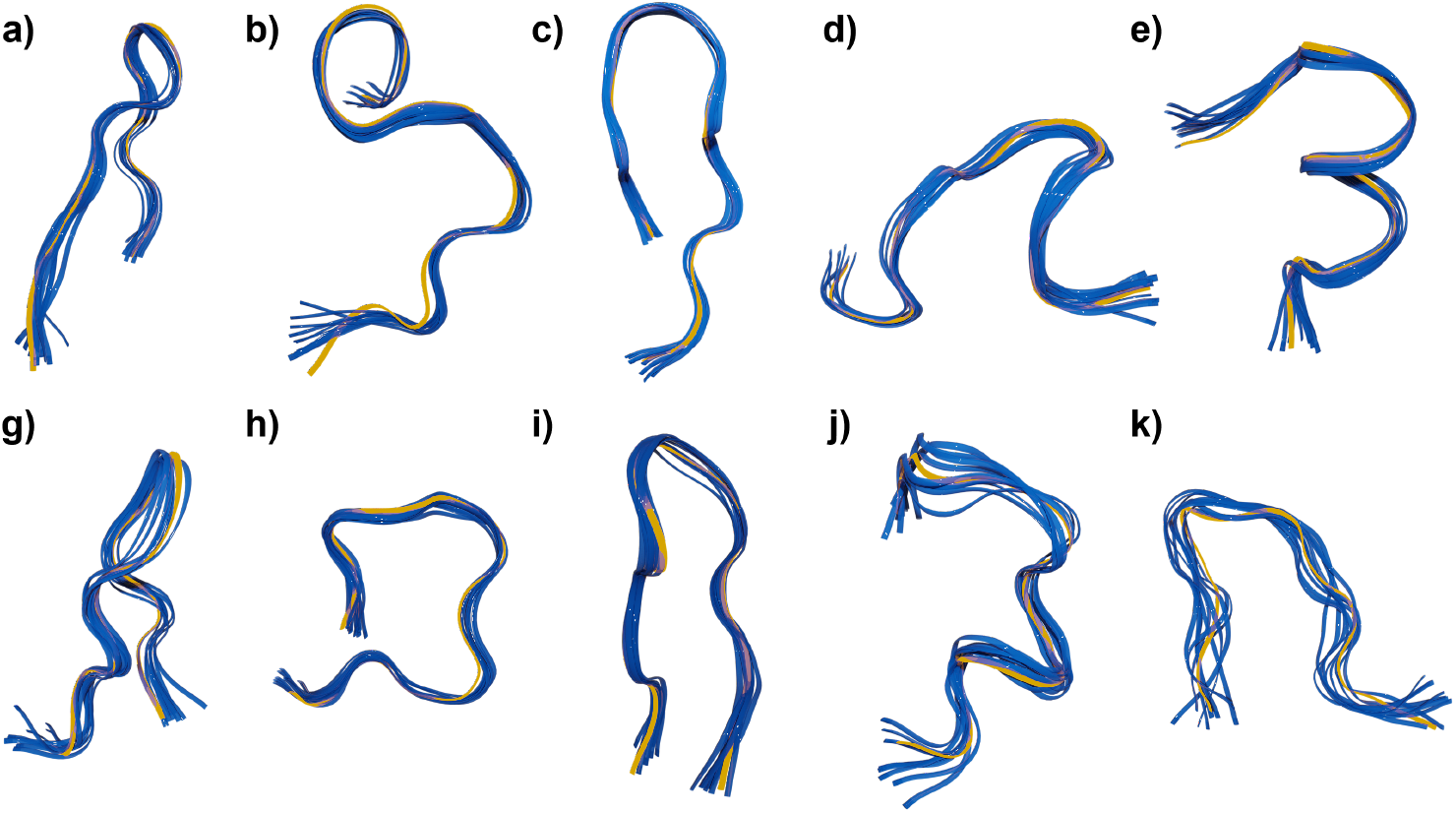
Overlaps of eQual clustering results at the maximum number of clusters (*k* =155 and MSD=1.7). Each conformation is colored in blue with the medoid colored in yellow.

**Figure 11.**
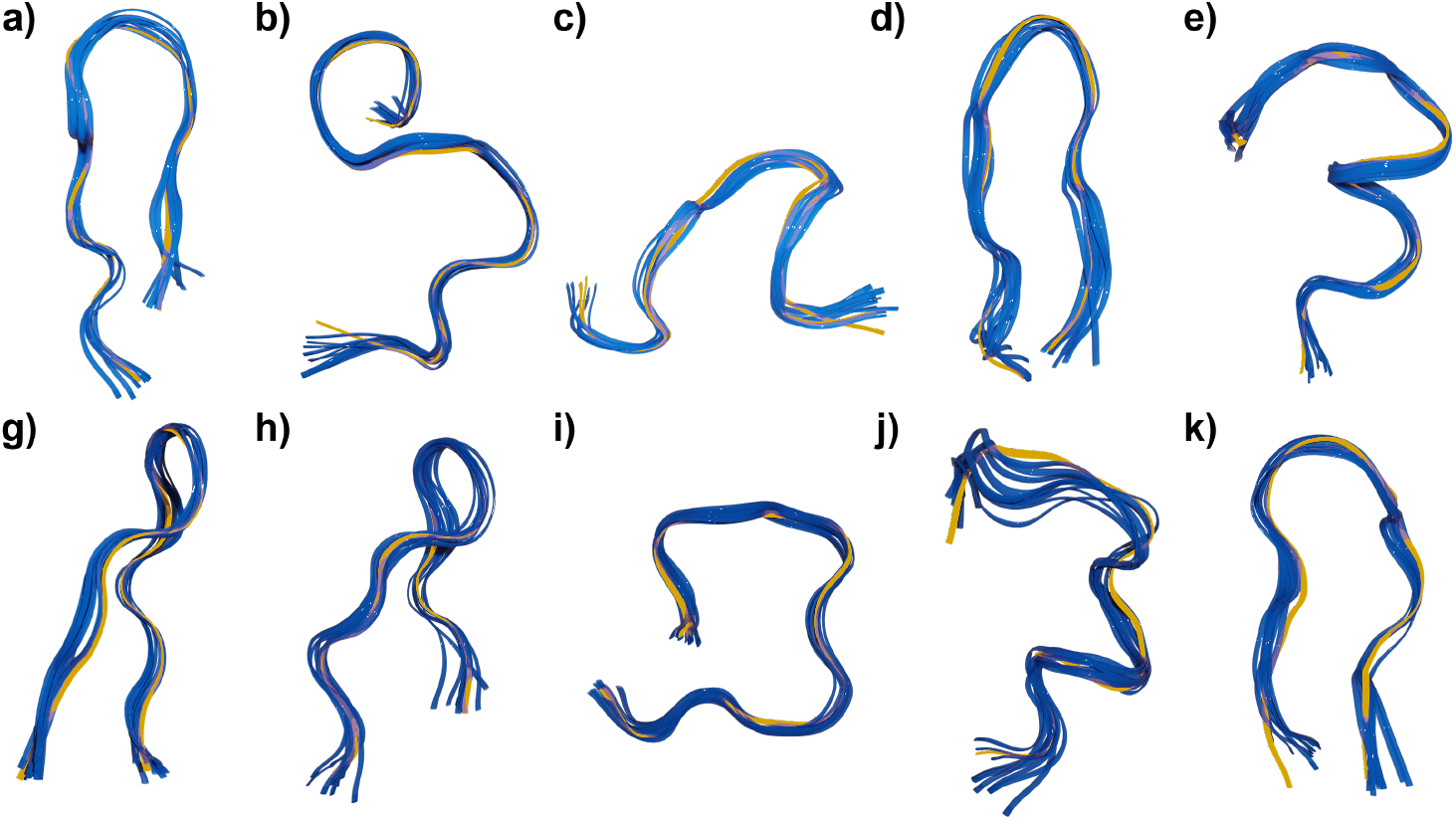
Overlaps of CADENCE clustering results when the number of outliers is 10 and MSD is 1.6. Each conformation is colored in blue with the medoid colored in yellow.

## Conclusion

This manuscript introduces CADENCE, a linear-scaling density-based clustering method that facilitates finding clusters with non-convex or arbitrary shapes. CADENCE builds upon eQual, a radial threshold clustering, which optimizes the selection of initial seeds, and all points within the radial threshold are accepted to the clusters. eQual clusters are, by construction, more tightly packed and mostly globular, while CADENCE clusters are more flexible and of varying shapes. Two CADENCE parameters were introduced, the number of outliers (n out) and the number of points (n points), and investigated to test its effects on the resulting number of clusters with the help of the CHI and DBI indicators. With (n out), the cluster behavior does not change drastically from eQual. Both eQual and n out have the same peak threshold with the maximum number of clusters, showing consistency between both methods, and their attractive ability of finding intrinsic thresholds in the data that guarantee maximum separability between the sets. With this extra flexibility of including two parameters, we found combinations resulting in higher CHI and lower DBI values, leading to better cluster quality when clusters are more irregular, something that is not possible with radial-like (e.g., eQual) or Voronoi-like (e.g., k-means). Furthermore, population metrics validated that the density expansion was indeed able to incorporate new conformations. Lastly, from the structural overlaps, the same motifs observed in eQual were also observed in CADENCE, which indicates that CADENCE is still able to maintain the tight partitioning of the data, while adding some new diversity to the clusters. In short, CADENCE provides a natural generalization of eQual (or other radial methods), showing how the clusters could be relaxed to adopt arbitrary shapes, without compromising their inner structure. CADENCE is publicly available as part of our MDANCE package:.

## Data and Software Availability

Clustering was performed with the open-source CADENCE module under the MDANCE software package that is available on GitHub https://github.com/mqcomplab/MDANCE. This includes the Molecular Dynamics input files and scripts to reproduce these results. Full documentation can be found in https://mdance.readthedocs.io.

## Acknowledgement

RAMQ and LC thank support from the National Institute of General Medical Sciences of the National Institutes of Health under award number R35GM150620.

